# Cultivation of Methanonezhaarchaeia, the third class of methanogens within the phylum Thermoproteota

**DOI:** 10.1101/2025.06.25.661132

**Authors:** Anthony J. Kohtz, Sylvia Nupp, Roland Hatzenpichler

## Abstract

Methane is a potent greenhouse gas that is largely produced through the activity of methanogenic archaea, which contribute to Earth’s dynamic climate and biogeochemical cycles. In the past decade, metagenomics revealed that lineages outside of the traditional Euryarchaeota superphylum encode genes for methanogenesis. This was recently confirmed through the cultivation of two classes of methanogenic Thermoproteota. Thus far, all methanogens within the Thermoproteota are predicted or were shown to be methylotrophic. The only exception to this is the Nezhaarchaea, which, based on metagenomics, are predicted to be CO_2_-reducing methanogens. Here, we demonstrate methanogenic activity in a third class of Thermoproteota, the Methanonezhaarchaeia. We expand the metabolic diversity of this class by cultivating a methylotrophic species, *Candidatus* Methanonezhaarchaeum fastidiosum YNP3N. We describe novel genes involved in methanogenesis that are not found in other methanogens. We investigate the metabolic diversity of Methanonezhaarchaeia, including metabolic modifications accompanying frequent loss of methanogenesis in this class. This highlights gaps in our understanding of the biochemistry, diversity, and evolution of non-traditional methanogens and their contributions to carbon cycling.

**Teaser:** The cultivation of a new group of methanogens illuminates their metabolic diversity and evolution of archaea.

## Introduction

Methanogenesis, the production of methane coupled to energy conservation, is a uniquely archaeal metabolism (*1, 2*). Through their activity, methanogens are responsible for mediating global methane emissions, which significantly affects Earth’s climate (*3*). While all methanogens share a unique methane-forming enzyme, methyl coenzyme-M reductase (MCR), different precursor molecules can be converted to methane via distinct pathways : (i) CO_2_-reducing, (ii) acetoclastic, (iii) methyl-dismutating, (iv) methyl-reducing, (v) methoxyl-dismutating, and (vi) alkanotrophic methanogenesis(*1*).

In addition to discoveries regarding their metabolism (*4-7*), our understanding of which archaea engage in methanogenesis has radically changed over the past decade. Traditionally, all methanogens were thought to belong to the superphylum Euryarchaeota. However, this view rapidly changed after several independent metagenomic studies discovered genes for anaerobic alkane-cycling in numerous uncultured archaeal lineages within the phylum Thermoproteota (*a*.*k*.*a*. TACK superphylum), including Korarchaeia (*5, 7*), Methanomethylicia (previously called Verstraetearchaeota) (*8*), Nezhaarchaea (*9*), Nitrososphaeria (*10, 11*), Aukarchaeles (*12*), and Bathyarchaeia (*13, 14*). In addition, even in traditional Euryarchaeota, multiple deep branching methanogenic and alkane degrading lineages have been discovered in the past decade, showing our understanding of the mediators of these metabolisms are not yet complete (*5, 6, 9, 15-18*). Only recently, the first three cultures of Korarchaeia and Methanomethylicia were recovered, enabling an experimental validation of their methanogenic physiology (*12, 19, 20*). The vast expansion in experimentally confirmed as well as potential non-traditional methanogens has wide-ranging implications for our understanding of global methane cycling, the evolution and biology of archaea, and the biotechnological potential of these novel methanogens (*1, 11, 21, 22*).

Almost all lineages of (potential) methanogens within the Thermoproteota are predicted (*5, 7-14*) or have been demonstrated (*12, 19, 20*) to be methyl-reducing methanogens. This idea is supported by the presence of methyltransferase genes and the lack of both the Wood-Ljungdahl Pathway (WLP) and the tetrahydromethanopterin S-methyltransferase (MTR) that can connect methanogenesis to the WLP. So far, the only lineage within the Thermoproteota that encodes both MCR and MTR are the ‘Nezhaarchaea’, which, based on metagenomics, are predicted to be CO_2_-reducing methanogens (*9, 11*). Nezhaarchaea were first discovered in hot spring samples (*9, 11*) but are also present in deep-sea hydrothermal systems (*23*), suggesting that they are important for methane cycling in high temperature environments. So far, the only experimental evidence supportive of a methanogenic lifestyle was a study demonstrating that Nezhaarchaea are the most abundant MCR-encoding archaea in some hot springs and transcribe MCR in methanogenic microcosms seeded with biomass from those hot springs (*24*). While this was an important step forward, most of their metabolism remains unresolved due to low recruitment of metatranscriptomic reads to genes beyond MCR and Nezhaarchaea remain enigmatic without cultures.

Here we demonstrate methylotrophic methanogenesis in the Nezhaarchaea through selective cultivation, fluorescence microscopy, and metagenomic and metatranscriptomic sequencing. We further assess the wider metabolic diversity in this lineage, suggesting loss of methanogenesis across this lineage is associated with rewiring of membrane-associated redox complexes.

## Results

### Cultivation of a new methylotrophic methanogen

We recently demonstrated the Lower Culex Basin (LCB) in Yellowstone National Park to be an area hosting a taxonomically diverse range of novel MCR-encoding archaea affiliating with the superphylum Thermoproteota (*25*). Notably, methanogenic cultures of the first two classes of Thermoproteotal methanogens, affiliating to the Korarchaeia and Methanomethylicia, were obtained from hot springs located in the LCB (*19, 20*). Furthermore, mesocosm experiments using sediments from hot spring ‘LCB003’ revealed that members of a third class of Thermoproteotal MCR-encoding archaea, Nezhaarchaea, became slightly enriched (up to 8% relative abundance) under methanogenic conditions when incubated with methanol under a N_2_/CO_2_ headspace (*25*).

To further explore the metabolism of Nezhaarchaea and attempt their cultivation, we incubated sediments from hot spring LCB003 in anoxic media under a variety of conditions designed to stimulate methylotrophic methanogenesis at the *in situ* temperature of 77 °C and pH of 6.5. We found that incubations with monomethylamine (MMA), dimethylamine, or trimethylamine, with or without isopropanol), and a N_2_/CO_2_ headspace repeatedly led to the enrichment of Nezhaarchaea along with pronounced methane production. Further incubations revealed that isopropanol is not required for enrichment and has little effect on total methane production (Fig. 1A). MMA exhibited the most consistent response in methane production, and all further work was performed using MMA as a substrate. The cultures exhibited long lag times of around 50-60 days when first inoculated into anoxic media, and this response was repeatable over multiple years of sampling from the same hot spring. Cultures also frequently stopped growing after 1-2 transfers into fresh media; however, we were able to eventually obtain a stable methanogenic culture for detailed study.

**Figure 1.**
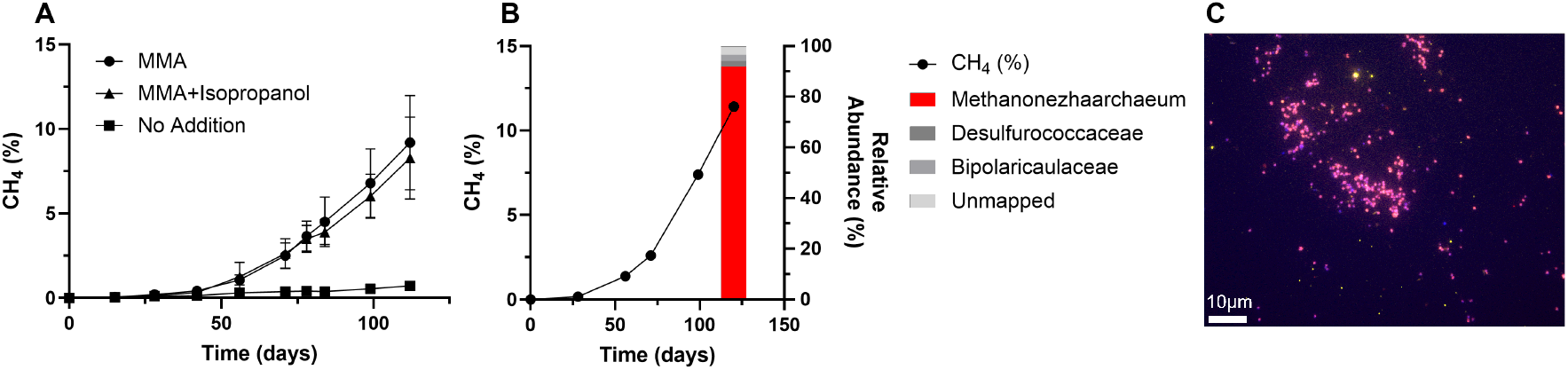
Enrichment cultivation and visualization of *Ca*. Methanonezhaarchaeum fastidiosum YNP3N. **(A)** Methane production by the culture under varying amendments: MMA only (n = 9), MMA and isopropanol (n = 9), and no addition (n = 3). Symbols are the mean from biological replicates and error bars are one standard deviation, where not visible the error bars are smaller than the symbol. **(B)** Long term cultivation on MMA and multiple transfers produced a sediment-free culture >90% enriched in strain YNP3N according to metagenomics (n = 1). **(C)** DOPE-FISH confirmed the high enrichment of YNP3. Abundant coccoid cells double hybridized (appearing in pink) with a general archaeal probe (Arch915, red) and a probe specific to Methanonezhaarchaea (Mnez277, yellow). Cells were counterstained with DAPI (blue). Bright yellow signals stem from auto-fluorescent sediment particles.

We found that incubations initiated with 50% H_2_ in the headspace did not develop any significant enrichment of Nezhaarchaea, suggesting that they may be adapted to low H_2_ concentrations or are outcompeted under those conditions. In contrast, the methanogenic korarchaeon *Ca*. Methanodesulfokora washburnensis, which we previously enriched from the same hot spring, rapidly increases in abundance under a 50% H_2_ headspace and methanol amendment (*19*). This indicates that these thermoproteotal methanogens inhabit different niches within the same hot spring sediment with regards to both substrate (methylamines vs. methanol) and hydrogen (no/low H_2_ vs. high H_2_) preferences.

Long read Nanopore and short read Illumina sequencing of the culture recovered a complete circular genome of the nezhaarchaeon, with a size of 1.75 Mbp and a GC content of 53% (Supplementary Table 1). Read mapping of the short read metagenome to the genome revealed that this archaeon dominated the culture at a relative abundance of 90.13% (Figure 1B, Supplementary Table 1). This strong enrichment was independently confirmed by DOPE-FISH (Figure 1C). In contrast, by mapping the raw reads of the LCB003 hot spring metagenome to the genome of the archaeon in culture, we estimate the relative abundance of the organism in its native habitat at 0.46%. Phylogenomic, average nucleotide identity (ANI), and GTDB-tk analyses showed that the cultured archaeon clusters within the Nezhaarchaea lineage and that it represents a new species, for which we propose the name *Candidatus* Methanonezhaarchaeum fastidiosum strain YNP3N (Figure 2A, Figure S1, Supplementary Table 1). We chose this name to emphasize the ‘methano-’ prefix typical in methanogen taxonomy, to preserve previously proposed names as much as possible (*i*.*e*., Nezhaarchaeota, Nezhaarchaea (*9, 11, 24*)), and to emphasize the fastidious and slow-growing nature of this organism. We note that this name change would necessitate slightly renaming this group using the type genus Methanonezhaarchaeum propagated up to class Methanonezhaarchaeia (see Etymology section at the end of the manuscript the and the Protologue in the Supplementary Online Information).

**Figure 2.**
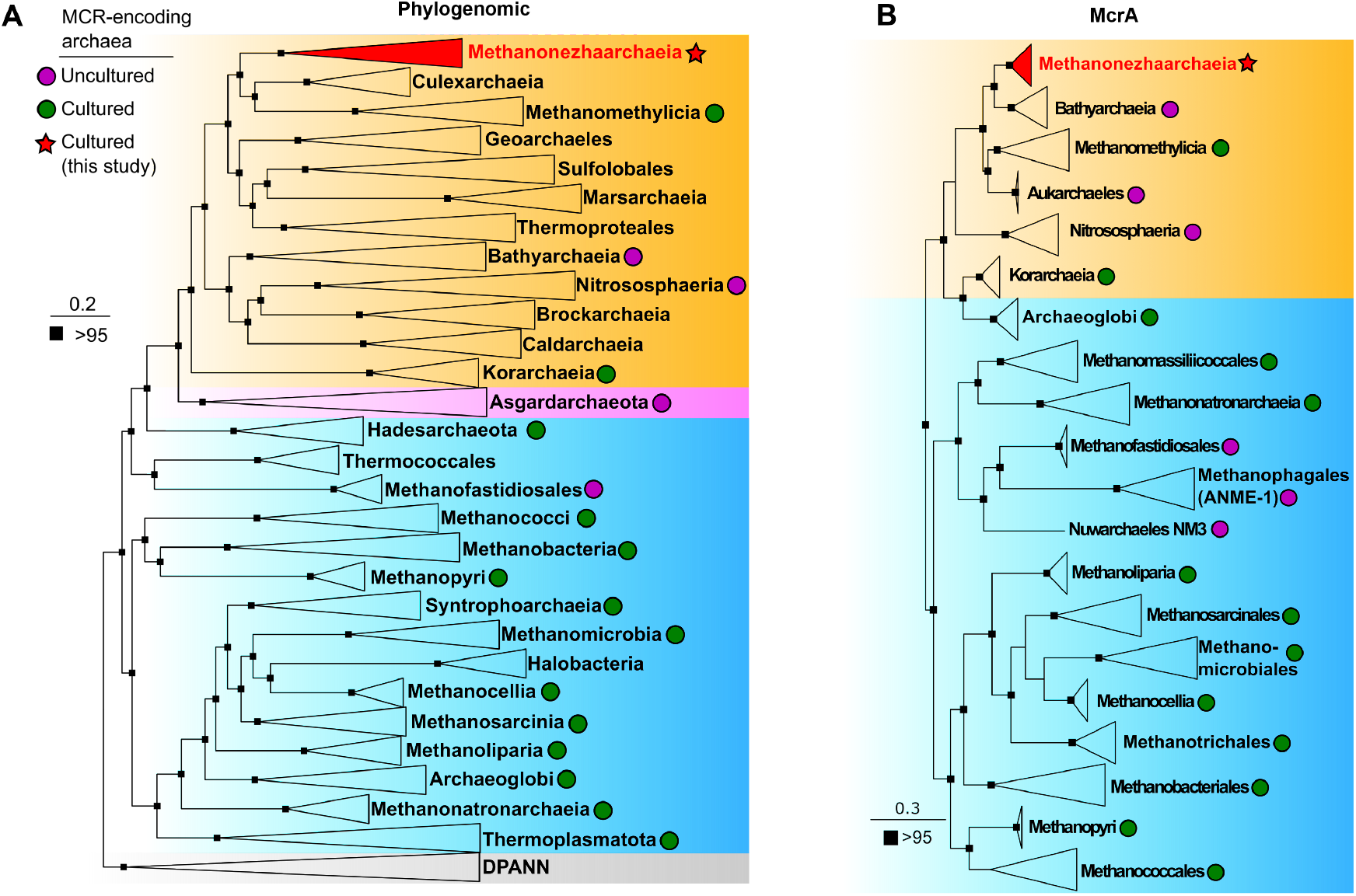
Phylogenetic placement of Methanonezhaarchaeia based on single copy marker genes and McrA. **(A)** Phylogenomic tree of archaea based on a set of 43 single-copy marker genes. Major groups of archaea are highlighted in orange (Thermoproteota/TACK superphylum), pink (Asgardarchaeota), blue (Euryarchaeota), and gray (DPANN). For inclusion, genomes were required to have at least 37 of the marker genes. Squares at the nodes indicate ultrafast bootstrap support, only values above 95 are shown. **(B)** Phylogenetic tree of McrA sequences demonstrating the affiliation of YNP3N within the Methanonezhaarchaeia (Thermoproteota, orange) compared to traditional Euryarchaeota (blue). Squares at the nodes indicate ultrafast bootstrap support, only values above 95 are shown.

Strain YNP3N encoded the only *mcrA* gene recovered in the metagenome assembly, and phylogenetic analysis of this gene placed it within the Methanonezhaarchaeia, with recently described Bathyarchaeia sequences as a sister group (Figure 2B) (*13*). We also found fragmented *mcrBCD* genes related to *Ca*. Methanoglobus hypatiae (*16*), known to metabolize MMA, in the metagenome assembly. However, no Methanoglobus *mcrAG* was recovered, the *mcrBCD* genes had low coverage, and mapping the metagenomic reads to the *Ca*. M. hypatiae genome revealed a relative abundance of 0.037%. Together, this suggests minimal impacts on the observed methane production by *Ca*. M. hypatiae. We also observed a second species of Methanonezhaarchaeia in the culture, with a low relative abundance of 1.6%. However, the MAG was non-circular, less complete (85.29%) and did not encode *mcr* genes (Supplementary Table 1).

A new FISH probe was designed to target Methanonezhaarchaeia. Application of this probe and a general archaeal probe revealed abundant coccoid cells that were labeled with both probes, confirming the high enrichment of strain YNP3N (Fig. 1C). Searches against public databases using the 16S rRNA and *mcrA* gene sequences showed that Methanonezhaarchaeia are globally prevalent in terrestrial hot springs and marine hydrothermal systems (Supplementary Table 2,3). However, it is important to note that Methanonezhaarchaeia likely avoided detection in the past because commonly used PCR primers exhibit multiple mismatches to their *mcrA* genes(*25*).

### Metabolism and gene expression

Reconstruction of the metabolic potential revealed that strain YNP3N encodes a more diverse metabolism for methanogenic substrates relative to previously published Methanonezhaarchaeia MAGs, which had been proposed to exclusively use H_2_/CO_2_ for methanogenesis. As expected, the MCR complex genes were highly expressed during methanogenic growth. During growth on monomethylamine, YNP3N highly expresses methyltransferase genes involved in the metabolism of this substrate (*mtmB, mtbC*) and expresses genes for using other methylamines (*mtbB, mttB*) at a lower level (Figure 3B). While there was not an associated *mtbA* gene, an *mtaA*-like gene is co-located and highly expressed, likely replacing the activity of *mtbA*, similar to the methyltransferase systems found in Methanomassillicoccales (*26*) and *Ca*. Methanglobus hypatiae (*16*). We also identified pyrrolysine biosynthesis genes, PylSBCD, and a Pyl-tRNA used to recode stop codons, notably found within the methylamine methyltransferase genes (*27-29*). Furthermore, we observed expression of diverse corrinoid transporter genes and cobalamin-binding proteins required for import of vitamin B_12_ and synthesis of cobalamin proteins found in methyltransferase systems (*30, 31*). This indicates the importance of this vitamin in the metabolism of this organism and suggests that in the native hot spring these transporters are likely important in uptake of B_12_ from the environment. Strain YNP3N also expressed both the tetrahydromethanopterin S-methyltransferase (MTR) complex and the methyl branch of the Wood-Ljungdahl Pathway (WLP), further indicating its capability for methyl-dismutating methanogenesis. This is in stark contrast to previously cultured Thermoproteota methanogens, which all lack a full MTR complex and most lack the WLP (*1, 10, 12, 19, 20*). We could not sustain growth of strain YNP3N without addition of MMA as a growth substrate, even with CO_2_/bicarbonate present, suggesting that YNP3N, contrary to its genomic potential, does not perform CO_2_-reducing methanogenesis *in vitro*. This finding highlights the importance of testing predictions from (meta)genomics through careful experiments and the indispensability of cultivation.

**Figure 3:**
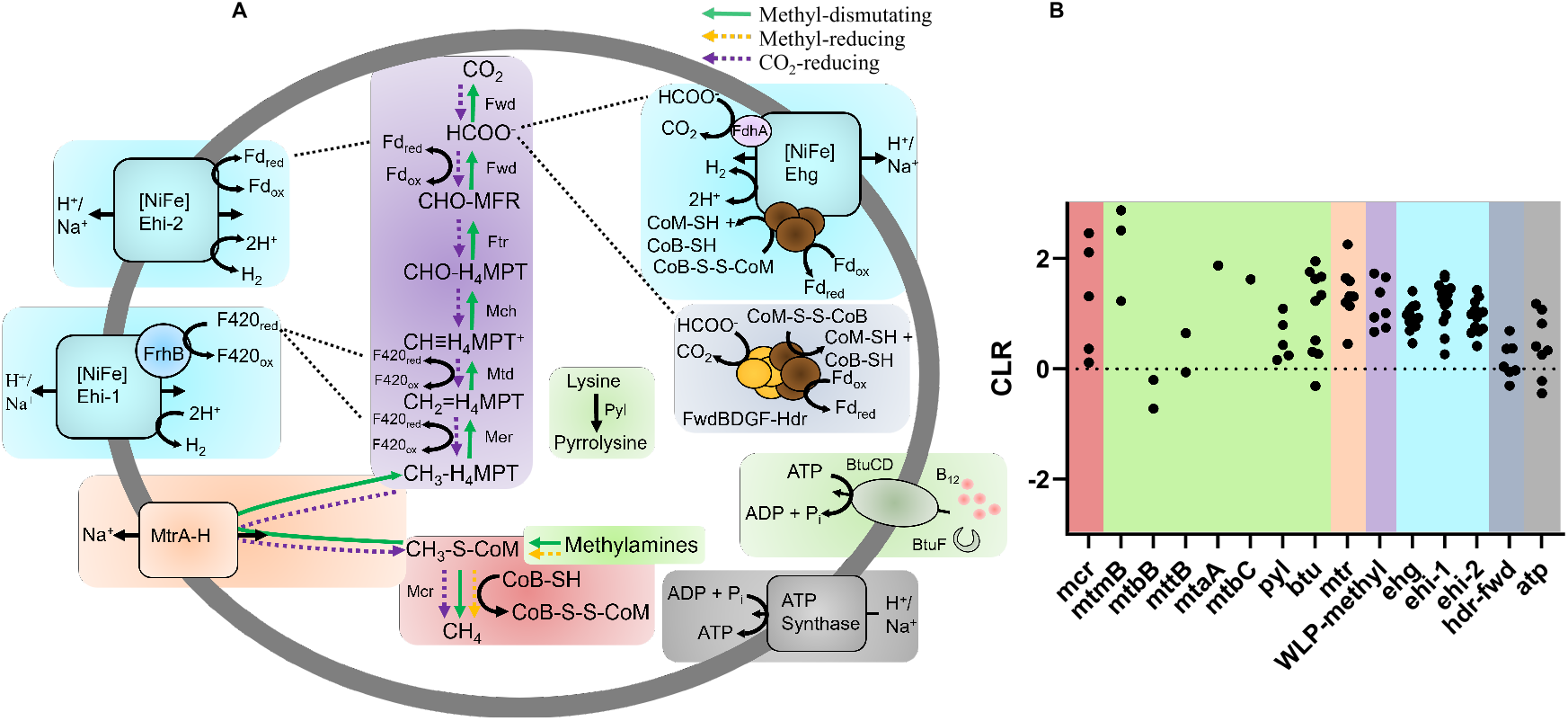
Proposed methane, hydrogen, and formate metabolism in *Ca*. M. fastidiosum YNP3N. **(A)** Metabolic reconstruction of methanogenesis, energy conservation, and vitamin B12 import in YNP3N. A list of genes used to construct this figure can be found in Supplementary Table 4. **(B)** Gene expression of YNP3N during growth on monomethylamine (n=1 due to low RNA yields) suggests a methyl-dismutating methanogenic lifestyle. Dotted line represents the geometric mean expression of all genes. Unless a specific subunit is listed, each dot represents the expression of a subunit in a multi-subunit enzyme or functional class, or expression of multiple copies of a particular gene. Explicit values can be found in Supplementary Table 5.

For heterodisulfide reduction, methanogens lacking cytochromes commonly have an electron-bifurcating cytoplasmic Mvh-Hdr complex that allows coupling the oxidation of H_2_ to the reduction of ferredoxin and heterodisulfide (*32*). In formate-utilizing methanogens, Mvh-Hdr can also form large complexes with FdhAB, to allow formate to substitute for H_2_ as an electron donor (*33*). YNP3N lacks FdhAB or MvhABG associated with a soluble HdrABC operon. Instead, a second gene cluster containing FwdBD subunits are found co-located with HdrABC in the YNP3N genome. The FwdBD subunits typically perform CO_2_-reduction to formate in hydrogenotrophic methanogens, with formate then migrating to the FwdA subunit (*34, 35*). Thus, this Fwd-Hdr complex may enable formate oxidation to be coupled with heterodisulfide and ferredoxin reduction. The additional co-located FwdF polyferredoxin and FwdG genes would likely allow to electronically couple the HdrABC and FwdBD subunits. This Fwd-Hdr organization is a new strategy distinct from the characterized Fmd-Hdr-Fdh or Fwd-Hdr-Mvh-Fdh complexes found in Methanosprillium or Methanococcus, respectively (*33, 36*).

Strain YNP3N also expresses a separate FwdABCD gene cluster co-located with other genes of the WLP, and here the FwdBD genes are fused, indicating gene duplication and fusion events enabled a repurposing of the second Fwd complex for involvement in heterodisulfide cycling.

Furthermore, for formate, hydrogen, and heterodisulfide cycling, YNP3N also expressed a membrane associated Group 4 [NiFe]-hydrogenase co-located with HdrABC and formate dehydrogenase (FdhA) subunits, termed Ehg (Fig. 3, Fig. S3) (*37*). Thus, this complex may be able to couple the oxidation of H_2_ or formate to the reduction of heterodisulfide and ion translocation for energy conservation. The FdhA subunit in YNP3N is similar in sequence (35-38% sequence identity) to those found in formate oxidizing methanogens within the Methanococcales, Methanobacteriales, and Methanocellales (*38-40*). However, with the presence of HdrA, this complex may also be capable of flavin based electron bifurcation and ferredoxin reduction. Internal formate cycling may take place during methyl oxidation through the WLP and may be sensitive to formate concentration. Attempts to grow strain YNP3N with MMA on a range of exogenous formate concentrations (2, 10, and 100 mM) were unsuccessful (not shown) and we were unable to identify a formate-specific transporter as found in other formate-utilizing methanogens (*38-40*).

Additionally, YNP3N encodes several other [NiFe] hydrogenases, suggesting that internal hydrogen cycling mechanisms could fuel methyl-dismutating methanogenesis. One expressed membrane bound group 4 [NiFe]-hydrogenase complex (Ehi-1, Fig. 3, Fig. S3) is flanked by *frhB*, suggesting this complex is involved in the oxidation of reduced F_420_ that could be produced by oxidizing methyl groups via the WLP during methyl-dismutating methanogenesis. Strain YNP3N encodes the biosynthetic pathway for producing F_420_, and we could not identify any other genes outside of *frhB* involved in recycling the F_420_ cofactor. A second expressed membrane bound group 4 [NiFe]-hydrogenase may deal with ferredoxin recycling produced by Fwd during methyl oxidation. This reduced ferredoxin could be oxidized by a second group 4 [NiFe] hydrogenase (Ehi-2, Fig. 3, Fig. S3) coupled with transport of ions across the membrane and the production of H_2_. Recycling of hydrogen produced through methyl oxidation may be recaptured by the Ehg hydrogenase complex, allowing for the direct coupling of heterodisulfide reduction and energy conservation.

The overall flexible metabolism of YNP3N suggests the use of multiple substrate and electron donor pools (methyl groups, H_2_, formate) may enable resilience during changing environmental conditions, as found in dynamic geothermal systems. Consistent with this, Methanonezhaarchaea have been observed to be stable populations and are often the dominant MCR-encoding organism in several hot spring systems (*24, 25*).

### Evolution and diversity of Methanonezhaarchaeia

Our surprising discovery of methylotrophic methanogenesis in the Methanonezhaarchaea, which previously had been predicted to only perform CO_2_-reducing methanogenesis, led us to reassess the wider metabolic diversity of this lineage. We observed 13 different species clusters, comprising three families within the Methanonezhaarchaeales, which we propose as ‘Methanonezhaarchaeceae’ (Mnzh.; f__WYZ-LMO8), Muzhaarchaeceae’ (Mza.; f B40-G2), and ‘Jinzhaarchaeceae’ (Jza.; f__JAWCJE01) (Figure 4A, Supplementary Table 5).

**Figure 4.**
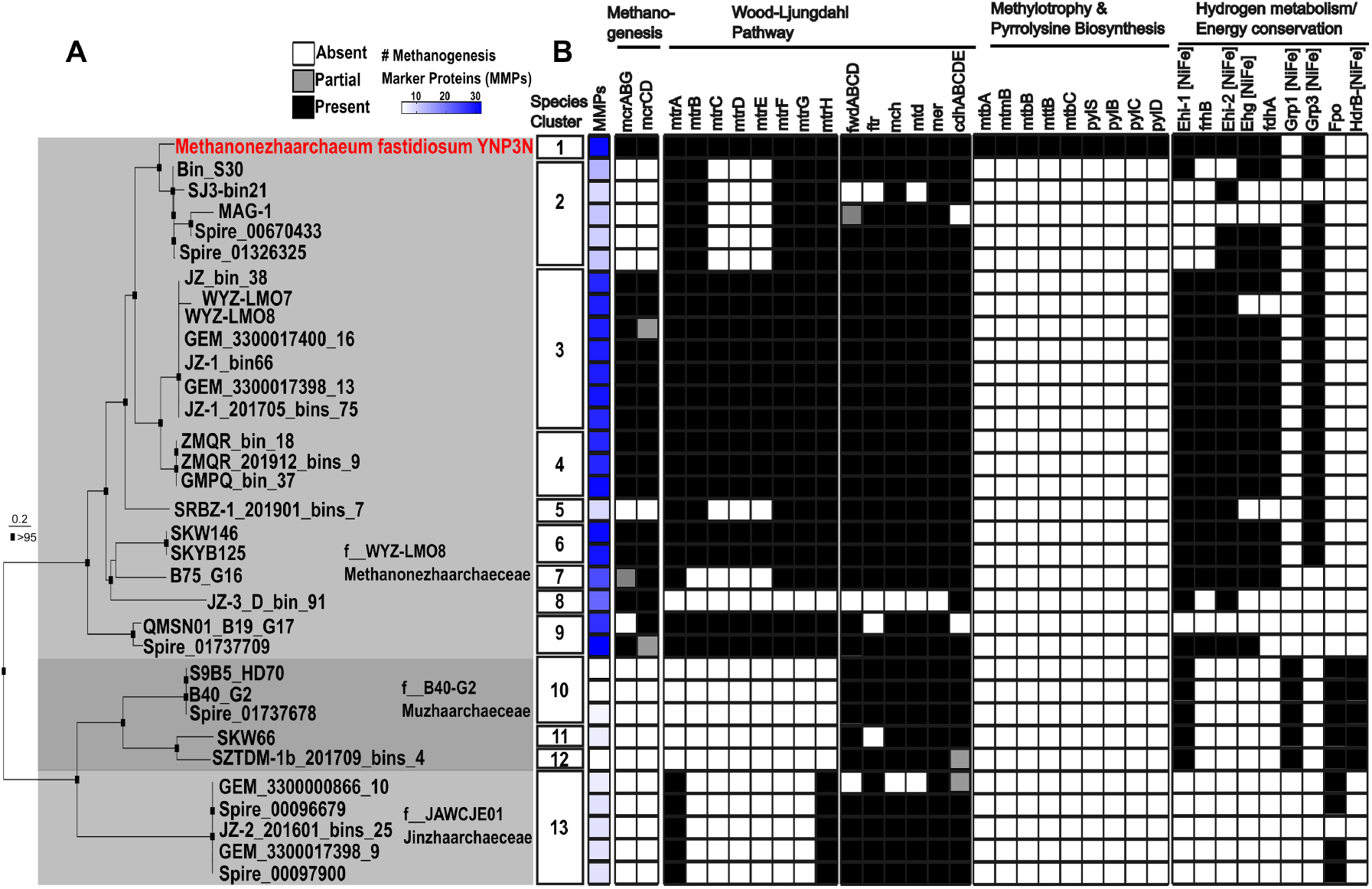
Comparative genomics of three families within Methanonezhaarchaeia. **(A)** Genome tree illustrating the diversity of Methanonezhaarchaeia. The phylogenetic tree shown here is a subset of the tree shown in Figure 2A. **(B)** Presence and absence of key metabolic genes and methanogenesis marker proteins.

*Ca*. M. fastidiosum YNP3N was the only genome found to contain genes for biosynthesis of pyrrolysine and methyltransferases for methylamines, indicating its unique capability for methylotrophic methanogenesis. We found that while nearly all Methanonezhaarchaeia species encode a complete or partial McrABGCD, all MAGs affiliated with its sister groups, Muzhaarchaeceae and Jinzhaarchaeceae, lacked these genes. This suggests a transition from a methanogenic to non-methanogenic lifestyle within the class Methanonezhaarchaeia. Consistent with this, all members of the Muzhaarchaeceae and Jinzhaarchaeceae, in addition to two species clusters within the Methanonezhaarchaeceae, encoded only a limited subset of the 38 methanogenesis marker proteins, including the one most closely related to strain YNP3N (87.4% ANI) (Figure 4B, Supplementary Table 6).

Furthermore, different patterns of MTR loss were observed, with Muzhaarchaeceae seemingly having lost this complex altogether, while members of the Jinzhaarchaeceae only encode the MtrA and MtrH subunits. Five of nine species clusters within the Methanonezhaarchaeceae encode a full MTR complex, while one lacked it completely, and three species clusters were missing MtrCDE. MtrCDE subunits interact in sodium ion pumping, and their loss could indicate an important intermediate stage in the loss of methanogenic potential, by uncoupling MTR from energy conservation in species that have already lost MCR (Figure 4) (*41*). Furthermore, the *mtrA* gene found in Jinzhaarchaeceae has a separate evolutionary history from those found in Methanonezhaarchaeceae, as it branches with sequences from the Asgardarchaeota while Methanonezhaarchaeceae sequences branch with Bathyarchaeia (Figure S4). Together, this is further evidence for a complicated history of loss and horizontal gene transfer of the MTR subunits in addition to repurposing of at least the MtrAH subunits in diverse non-methanogenic lineages (*21, 22, 42*).

Compared to the pattern observed for MTR, the methyl and carbonyl branches of the WLP were largely conserved across all three families. This is consistent with the highly flexible nature of this pathway that is connected to methanogenesis through MTR, but also serves multiple separate catabolic and anabolic roles in methanogenic and non-methanogenic microbes, including the biosynthesis of purines, thymidylate, methione, and acetyl-CoA (*43-45*).

For Methanonezhaarchaeceae, the membrane associated energy conservation and redox modules are largely conserved across the 8 identified species clusters, with numerous group 4 [NiFe] hydrogenases, including the HdrB and FdhA associated Ehg complex and the FrhB-associated Ehi-2 complex (Figures 3, 4). For Muzhaarchaeceae, we observed a large remodeling of membrane associated complexes, with a group 1 [NiFe] hydrogenase, Fpo, and a group 4 HdrB-associated [NiFe] hydrogenase similar to the Ehd complex found in the Methanomethylicia and Culexarchaeia (Figure 4, Figure S3) (*5, 37*). In contrast, Jinzhaarchaeceae exhibit a more minimalistic electron transport chain, encoding a single Fpo-like complex. Further cultivation and high-quality genomes of diverse Methanonezhaarchaeia are required to further untangle the stepwise modifications of MTR, hydrogenases, and Fpo-like complexes that lead to a loss of methanogenic potential and the repurposing of the enzymatic remnants of methanogenesis into a new lifestyle.

## Conclusion

Together, our results confirm methanogenesis in a third lineage of Thermoproteota, the Methanonezhaarchaeia. We expand the metabolic diversity of this group by cultivating a methylotrophic species from a group that hitherto had been assumed to only perform CO_2_-reducing methanogenesis based on metagenomic predictions. These findings highlight the need for continued sampling of non-traditional methanogen genomes and cultures, which likely harbor further yet-unidentified diversity in their methanogenic pathways, and to test (meta)genomic prediction by experimentation. Multiple novel gene arrangements of methanogenesis genes and associated redox and energy conserving complexes were identified in diverse Methanonezhaarchaeia genomes and found to be expressed in the YNP-3N culture. This continues the trend of novel discoveries in methanogenic members of the Thermoproteota and identifies gaps in our understanding of the biochemistry and diversity of methanogens.

Methanogenesis was independently lost multiple times in archaea, leading to a patchy distribution of this metabolism (*5, 21, 22*). We further find that methanogenesis likely is not a feature of all members of the Methanonezhaarchaeia, and that this metabolic transition was driven by loss of methanogenesis genes (MCR, MTR) and remodeling of membrane associated energy conservation and redox modules. Obtaining more diverse cultures will aid in resolving the full metabolic diversity of these non-traditional methanogens and their close, yet uncultured, non-methanogenic relatives.

## Etymology

YNP3N is a methanogen affiliated with the phylum Thermoproteota (*9*) for which we propose the candidate status Methanonezhaarchaeum fastidiosum sp. nov. For a protologue, please see the supplementary online information.

Etymology: Me.tha.no.ne.zha.ar.chae’eum. N.L. pref. methano-pertaining to methane metabolism; N.L. neut. pl. n. archaeum, “ancient”, referring to the cell’s affiliation to the archaeal domain; N.L. neut. n. Methanonezhaarchaeum, methanogen named after Nezha, an immortal in Chinese mythology (*9*). fas.ti.di.o’sum L. neut, adj. fastidiosum, referring to the fastidious growth of this organism.

Locality: Sediment from a hot spring, identified as feature LCB003 (*25*).

Diagnosis: a thermophilic methyl-dismutating methanogen of the phylum Thermoproteota that grows as coccoid cells.

## Methods

### Source of inoculum and cultivation

Sediment was retrieved from hot spring LCB003 (44.57763, −110.78957; November 2022; 77 °C; pH 6.5) as previously described and stored at room temperature (*19, 25*). Enrichment cultures were established using anoxic medium as previously described (*19, 20*), with minor modifications. The medium had a pH of 6.5, containing a base of (per liter): KH_2_PO_4_, 0.5 g; MgSO_4_·7H_2_O, 0.4 g; NaCl, 0.5 g; NH_4_Cl, 0.4 g; CaCl_2_·2H_2_O, 0.05 g; 2-N-morpholino-ethanesulfonic acid (MES), 2.17 g; yeast extract, 0.1 g; and 0.002% (w/v) (NH_4_)_2_Fe(SO_4_)_2_·6H_2_O, 5 mM NaHCO_3_, 1 mL trace element solution SL-10, 1 mL Selenite-Tungstate solution, 1 mL CCM vitamins, 0.0005% (w/v) resazurin, 10 mg of coenzyme-M, 2 mg sodium dithionite, 1 mM dithiothreitol and 1 mM Na_2_S·9H_2_O. Sediment slurry was dispensed into serum bottles (10% v/v) and then sealed with thick butyl-rubber stoppers and aluminum crimps in an anoxic glove bag (Coy). The headspace of the enrichment was exchanged with N_2_:CO_2_ (90:10) gas for 5 minutes and set to 200 kPa. Monomethylamine (MMA) was added to a final concentration of 10 mM. Cultures were incubated at 77 °C upright in the dark and without shaking. While attempting to optimize our media and obtain a stable culture, we made the following modifications for the final media: (i) in the original incubation isopropanol was also added at a concentration of 10 mM, but was subsequently omitted from the medium, because it did not affect the rate of growth or final methane concentration (ii) replaced MgSO_4_ with MgCl_2_·6H_2_O at 0.33 g (per liter), and (iii) removed choline from the CCM vitamins. Transfers of these batch cultures were made with 20% v/v inoculum into fresh medium when methane concentrations reached 8-12%. Dimethylamine (DMA) and trimethylamine (TMA) were also tested for methane production with and without isopropanol. However, these higher amines led to more inconsistent methane production profiles relative to MMA, and we focused on MMA.

### DNA extraction and metagenome sequencing

Under anoxic conditions, 10 mL of the YNP3N culture was collected aseptically with a needle and syringe and concentrated on a sterile 0.2-micron polycarbonate membrane filter (Nuclepore) with vacuum filtration. The filter was then immediately placed into a Zymo BashingBead tube containing 750 µL of ZymoBIOMICS lysis solution. DNA was extracted using a ZymoBIOMICS DNA miniprep kit following the manufacturer’s instructions. Illumina short-read library construction was performed with an Illumina DNA Prep kit following the manufacturer’s instructions and was sequenced on an Illumina NovaSeq X Plus platform with 2 x 151 bp paired-end read chemistry performed at SeqCenter (Pittsburgh, PA, USA). Nanopore long-read sequencing was performed in-lab using a MinION platform and a R10.4.1 flow cell (Oxford Nanopore Technologies). Library preparation was performed using a rapid barcoding kit (SQK-RBK114.24) following the manufacturer’s instructions and the flow cell was run for 27 hours. Base-calling was performed with dorado (v.0.7.1) using model sup@v.5.0.0. The resulting BAM file was demultiplexed using dorado demux (--emit-fastq).

### Metagenome assembly, polishing, binning, and quality assessment

An initial Nanopore assembly was performed with metaFlye (*46*) (v.2.9.4), which produced a circular assembly of the strain YNP3N genome. This genome was further polished using Illumina short reads using one round of Polypolish (v.0.6.0) followed by PyPolca (v.0.3.1; --careful) as recently suggested, given the high sequence coverage of the YNP3N genome (*47, 48*).

A short-read assembly was also generated for analysis of co-enriched microbes, with raw metagenomic reads with Illumina read quality, linker and adaptor trimming, artefact and common contaminant removal performed using the rqcfilter2 pipeline (maxns=3, maq=20) and error correction with bbcms (mincount=2, hcf=0.6) from the BBTools suite v.38.94 (Bushnell B. 2014. BBMap: a fast, accurate, splice-aware aligner. https://sourceforge.net/projects/bbmap). Resulting reads were then assembled with megahit (v.1.2.9; --k-list 27,37,47,57,67,77,87,97,107,117,127,141 –min-count 1) (*49*). Coverage was determined by mapping reads to the assembly using minimap2 (v.2.26-r1175.; -x sr) (*50*). Contigs with a length of 2kb or greater were used for binning with SemiBin2 (v.2.0.2). CheckM (v.1.1.3) was used to assess completeness and redundancy of the resulting MAGs (*51*). GTDB-tk (v.2.4.0) with the R220 release was used for assigning initial taxonomy (*52*). CoverM (v.0.7.0) with default settings was used to generate relative abundances of each MAG (*53*). CoverM was also used to estimate the relative abundance of the YNP3N genome in hot spring LCB003 using quality filtered reads from a previous sediment metagenome (IMG Genome ID 3300028675) (*19*).

### Annotation and reconstruction of metabolic potential

MAGs were annotated with Prokka (v.1.14.6), Interproscan (v.5.70-102.0) (*54, 55*). Annotation refinement and confirmation was done by submission of protein sequences to the NCBI conserved domain database and HHpred using default settings. Sequences of [NiFe] hydrogenases (PF00374) were checked for conserved CXXC motifs at the N and C terminus and submitted to HydDB for initial classification (*56*). Hydrogenase classification was further refined with phylogenetic analysis (below).

### RNA extraction, sequencing, and analysis

Two replicate cultures were grown on 10 mM monomethylamine under a N_2_/CO_2_ headspace. RNA was extracted from these cultures during active methane production. Culture bottles were cooled from 77 °C by placing them in an ice bath that was then placed at -20 °C for 15 minutes, and vials were subsequently kept on ice during the following procedure. Cells were rapidly concentrated onto sterivex filters, which were then cut open and placed into a Zymo BashingBead lysis tube (Zymo Research) containing 800 µL of TRIzol (Invitrogen) and vortexed at maximum speed for 20 minutes. The tubes were centrifuged at 14,000x g for 15 minutes and the supernatant was used for RNA purification with the Zymo Direct-Zol kit according to the manufacturer’s instructions and including the DNAse I step. RNA extract from the replicate cultures was pooled, and purified RNA was used for library preparation and sequencing at SeqCenter (Pittsburgh, PA, USA). RNA was treated again with DNAse I (Invitrogen) and library preparation was performed with Illumina’s Stranded Total RNA Prep Ligation with Ribo-Zero Plus kit and 10 bp unique dual indices. The library was sequenced on a NovaSeq X Plus platform, producing paired 2 x 150 bp reads. Demultiplexing, quality control, and adaptor trimming was performed with bcl-conver (v.4.1.5). Reads were further processed with rqcfilter2 (rna=t, trimrnaadapter=t, qtrim=rl, trimq=10, maq=20, maxns=0, minlen=50, mlf=0.33, removeribo=f) and mapped to reference genomes from the YNP3N culture using salmon (v.1.10.2; --validateMappings --seqBias) (*57*). Reads mapping to rRNA genes were excluded before normalization. Reads mapping to each gene were normalized to gene length before calculating the center log ratio (CLR; log_10_).

### Phylogenetic analysis

A set of 43 single-copy marker proteins was collected from Nezhaarchaeia MAGs and reference archaeal genomes as previously described (*20, 37*). Reference Nezhaarchaeia MAGs were collected from public databases and recent publications and only used if they were estimated at >75% completeness and had at least 37 out of 43 phylogenomic marker proteins (*58-60*). These markers were aligned with MAFFT-linsi, trimmed with trimAL (-gt 0.7) and concatenated to produce a final alignment of 9,913 positions (*61, 62*). IQtree2 was used to reconstruct a maximum-likelihood phylogenetic tree using the best fit LG+F+R10 model, and 1,000 ultrafast bootstraps (*63*).

The YNP3N McrA protein sequence was extracted and aligned with reference archaeal McrA sequences with MAFFT-linsi, trimmed with trimAL (-gt 0.5), producing a final alignment of 556 positions. IQtree2 was used to reconstruct a maximum-likelihood phylogenetic tree using the LG+C60+F+G model and 1,000 ultrafast bootstraps.

Reference archaeal 16S rRNA sequences ≥1000bp were aligned with the YNP3N 16S rRNA sequence using MAFFT-linsi and trimmed with trimAL (-automated1), producing an alignment of 1,498 positions. IQtree2 was used to reconstruct a maximum-likelihood phylogenetic tree using the best-fit SYM+R8 model and 1,000 ultrafast bootstraps.

Reference MtrA sequences were collected from diverse methanogenic and alkane degrading lineages and aligned with the YNP3N sequence using MAFFT-linsi and trimmed with trimAL (-automated1), producing an alignment of 242 positions. IQtree2 was used to reconstruct a maximum-likelihood phylogenetic tree using the best-fit LG+F+R6 model and 1,000 ultrafast bootstraps.

Reference sequences of catalytic subunits of group 1, 2, 3, and 4 [NiFe] hydrogenases (PF00374) and Fpo-like subunits (PF00346) were collected from HydDB and previous studies and aligned with sequences collected from Methanonezhaarchaeia genomes (*20, 56*). Sequences were aligned using MAFFT-linsi and trimmed with trimAL (-gt 0.5), producing an alignment of 423 positions. IQtree2 was used to reconstruct a maximum-likelihood phylogenetic tree using the LG+C60+F+G model and 1,000 ultrafast bootstraps.

### Fluorescence *in situ* hybridization

Subsamples of the YNP3N culture were fixed using 2% paraformaldehyde for 1 hour at room temperature as previously described (*20*). An oligonucleotide probe targeting most 16S rRNA sequences of Nezhaarchaea (equivalent to Terrestrial Miscellaneous Crenarchaeal Group (TMCG) in Silva release 138) was designed using the probe design tool in ARB (Mnez277, 5’-ACGGCCCGTACCCGTTATCG-3’). DOPE-FISH was performed as previously described (*20*) using the Mnez probe doubly labelled with Cy3 fluorophores (obtained from IDT-DNA) and a hybridization time of 3 hours at 10% formamide. Mnez277 was used in combination with the general archaeal probe Arch915, which was doubly labeled with Alexa Fluor 647 and cells were counterstained with the DNA stain 4′,6-diamidino-2-phenylindole (DAPI). Negative controls used double-labeled NON-EUB338 performed in parallel.

## Acknowledgments

We thank the US National Park Service for permitting work in YNP under permit number YELL-SCI-8010.

## Funding

U.S. Department of Energy, Office of Science, Biological and Environmental Research DE-SC0025661 (RH)

U.S. National Science Foundation, Biological Oceanography OCE-2049445 (RH)

Simons and the Gordon and Betty Moore Foundations 737750 (RH)

## Author contributions

Conceptualization: AJK, RH

Methodology: AJK, SN

Investigation: AJK, SN

Visualization: AJK

Supervision: RH

Writing - original draft: AJK

Writing - review & editing: AJK, SN, RH

## Data and materials availability

Metagenomic and metatranscriptomic data were deposited on NCBI under the Project ID PRJNA1272160.

