## Supplementary material for "Cultivation of Methanonezhaarchaeia, the third class of methanogens within the phylum Thermoproteota": SI text and figures

**This PDF file includes:**

Protologue
Figs. S1 to S4

**Other Supplementary Materials for this manuscript include the following:**
Tables S1 to S6 (xls format)

**Protologue**

**Class**

*Candidatus* Methanonezhaarchaeia class nov.

Me.tha.no.ne.zha.ar.chae'ia. N.L. neut. n. Methanonezhaarchaeales, type order of the class. L. -ia, ending to designate a class; N.L. fem. pl. n. Methanonezhaarchaeia, the Methanonezhaarchaeum class. The description is the same as for Methanonezhaarchaeum gen. nov.

**Order**

*Candidatus* Methanonezhaarchaeales order nov.

Me.tha.no.ne.zha.ar.chae'ales. N.L. neut. n. Methanonezhaarchaeceae, type family of the order. L. -ales, ending to designate an order; Methanonezhaarchaeales, the Methanonezhaarchaeum order. The description is the same as for Methanonezhaarchaeum gen. nov.

**Family**

*Candidatus* Methanonezhaarchaeceae fam. nov.

Me.tha.no.ne.zha.ar.chae'ceae. N.L. neut. n. Methanonezhaarchaeum, type genus of the family. L. -ceae, ending to designate a family; Methanonezhaarchaeaceae, the Methanonezhaarchaeum family. The description is the same as for Methanonezhaarchaeum gen. nov.

**Genus**

*Candidatus* Methanonezhaarchaeum gen. nov.

Me.tha.no.ne.zha.ar.chae'eum. N.L. pref. methano- pertaining to methane metabolism; N.L. neut. n. archaeum, "ancient", identifying this cell as a member of the domain Archaea; N.L. neut. n. Methanonezhaarchaeum, methanogenic archaeon named after Nezha, an immortal in Chinese mythology[1]. The type species is Candidatus Methanonezhaarchaeum fastidiosum sp. nov.

**Species**

*Candidatus* Methanonezhaarchaeum fastidiosum sp. nov.

fas.ti.di.o'sum L. neut, adj. fastidiosum, referring to the fastidious growth of this organism. This archaeon was cultured from an unnamed hot spring in Yellowstone National Park that we identified as feature LCB003 in a recent survey[2]. The organism is an obligate anaerobic thermophile that performs methyl-dismutating methanogenesis and grows as coccoid cells.

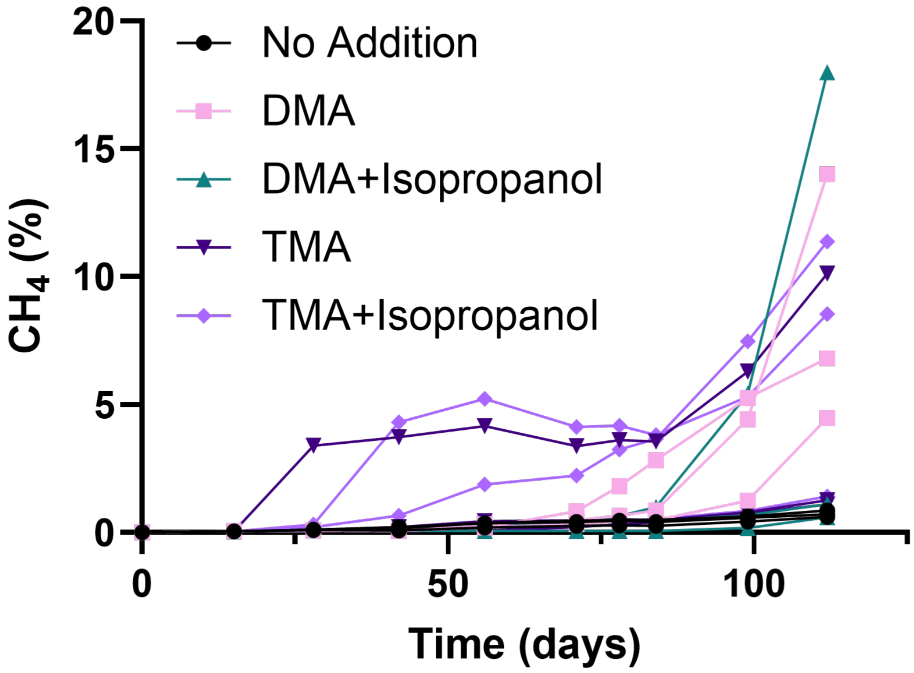


**Fig. S1. Methane production in incubations with dimethylamine (DMA) and trimethylamine (TMA), with or without isopropanol.** Each individual biological replicate is shown (n=3 for each condition). Methylamines and isopropanol were added at 10 mM final concentration.

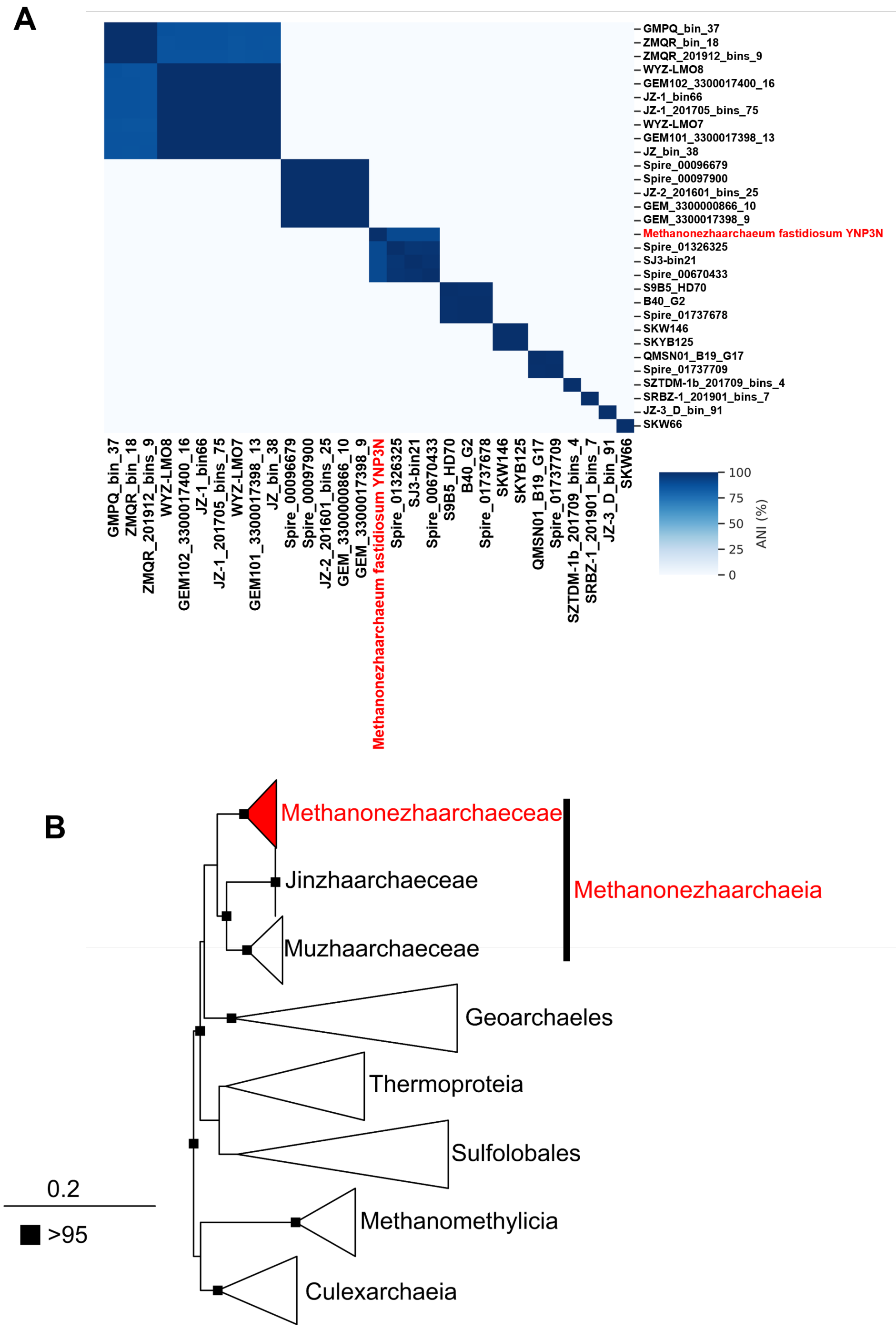


**Fig. S2. Average Nucleotide Identity (ANI) and phylogenetic analysis of 16S rRNA. (A)** ANI clusters across the Methanonezhaarchaeia showing placement of YNP3N as a separate species cluster. **(B)** Phylogenetic tree of the 16S rRNA showing YNP3N clustering within the Methanonezhaarchaeia. Squares at the nodes indicate ultrafast bootstrap support, only values above 95 are shown.

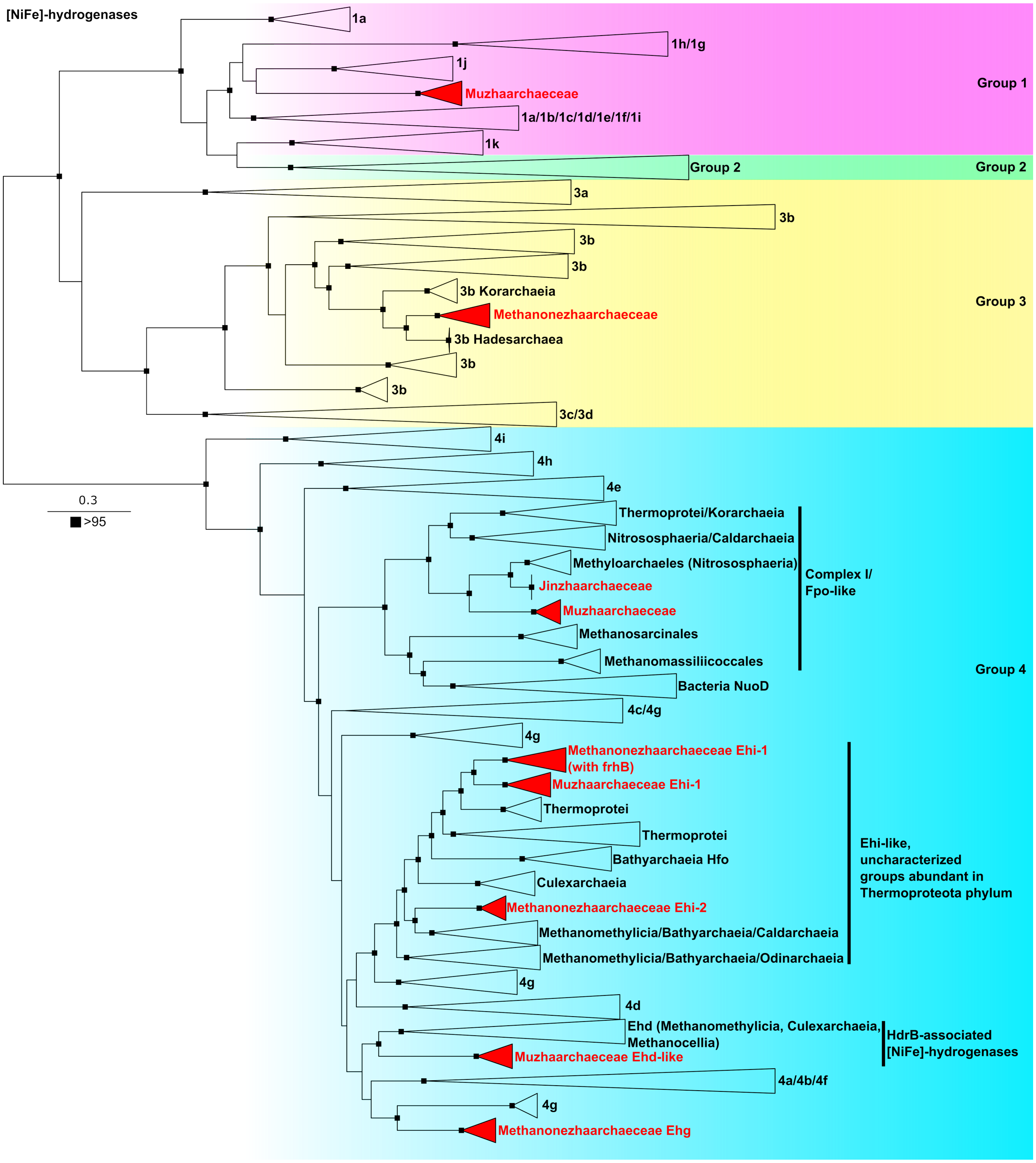


**Fig. S3. Phylogenetic tree of group 1, 2, 3, and 4 [NiFe] hydrogenases found in Methanonezhaarchaeia genomes (red).** The phylogenetic tree was constructed from an alignment of 423 positions and the LG+C60+F+G model. Squares at the nodes indicate ultrafast bootstrap support, only values above 95 are shown.

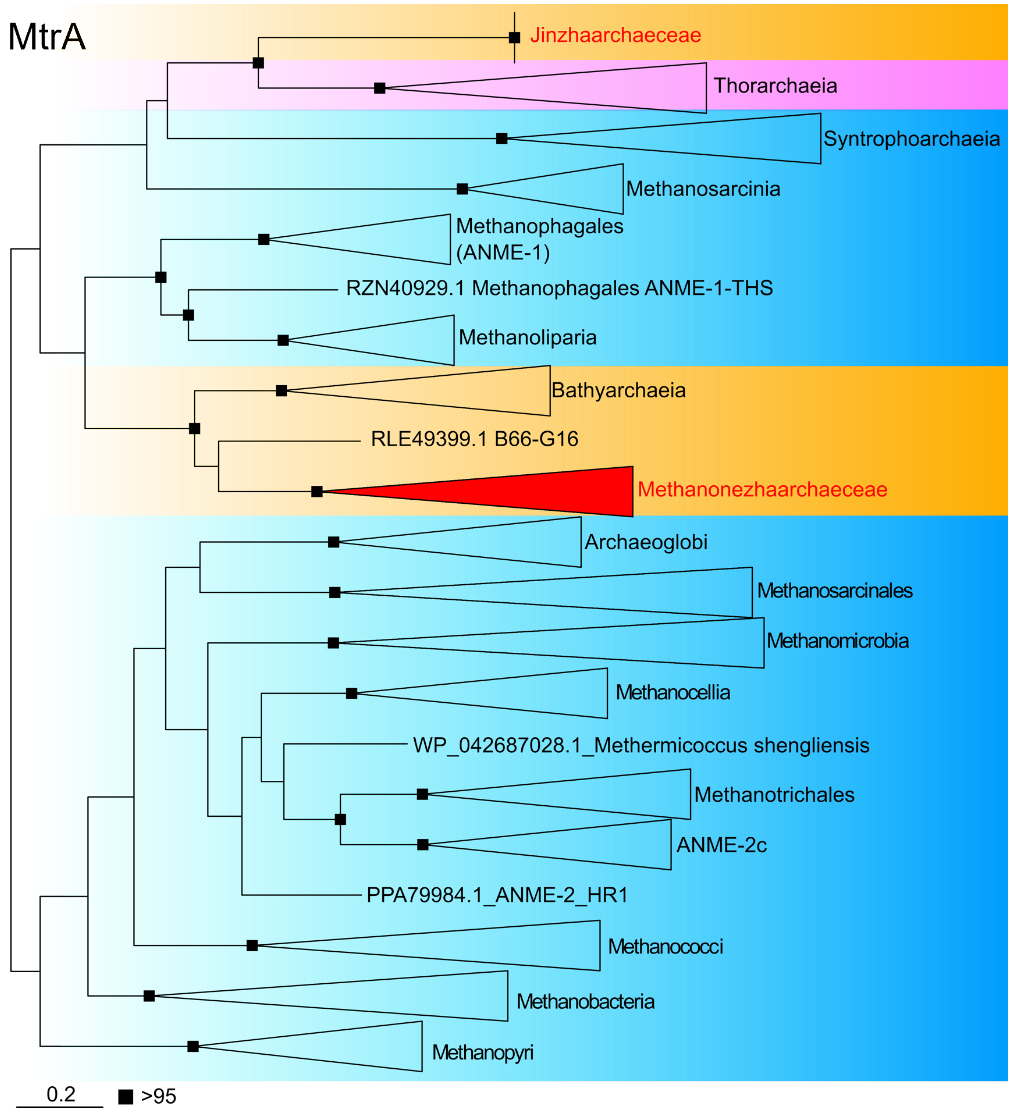


**Fig. S4. Phylogenetic tree of MtrA sequences collected from reference archaeal genomes and Methanonezhaarchaeia.** Two separate lineages are found in Methanonezhaarchaecea and Jinzhaarchaecea (red). The phylogenetic tree was constructed from an alignment of 242 positions, the best fit LG+F+R6 model, and 1,000 ultrafast bootstraps.

**Table S1 (separate sheet in xls file)**

Summary of the Ca. M. fastidiosum YNP3N genome, and Metagenome Assembled Genomes (MAGs) recovered from the enrichment metagenome.

**Table S2 (separate sheet in xls file)**

Distribution of 16S rRNA genes affiliated with Methanonezhaarchaeum in NCBI and IMG databases.

**Table S3 (separate sheet in xls file)**

Distribution of mcrA genes affiliated with Methanonezhaarchaeum in IMG databases.

**Table S4 (separate sheet in xls file)**

Summary of metatranscriptomic data shown in Figure 3B.

**Table S5 (separate sheet in xls file)**

Reference genomes of Methanonezhaarchaeia.

**Table S6 (separate sheet in xls file)**

Presence and absence of methanogenesis marker proteins in Methanonezhaarchaeia MAGs.
